# Auditory brainstem response amplitude predicts age-related forward masking temporal-processing deficits in normal-hearing listeners

**DOI:** 10.64898/2026.09.21.753073

**Authors:** Chengjie G. Huang, Milena Costantino, Roksana Soleimanpour, Taylor Beinke, Samira Anderson, Matthew J. Goupell

## Abstract

Aging has profound effects on sensory systems, such that older adults can report problems in hearing despite having normal audiometric thresholds. While it is known that older listeners have poorer temporal processing than younger listeners, there is limited understanding of how these deficits interact with effects of sound level, in particular when they are above conversational level (e.g., > 65 dB SPL). Here we wanted to investigate the link between temporal processing and sound level as a function of age in normal-hearing listeners. Using a forward-masking paradigm, auditory brainstem responses (ABRs) and behavioral detection thresholds were measured at different masker levels. It was found that with increasing age and masker sound level, older listeners’ Wave V amplitudes decreased and directly correlated with behavioral detection thresholds of the probe stimulus in the forward-masking paradigm. These results elucidate the fundamental effects of aging on temporal processing and its interactions with sound level in the central auditory system. The results of this study have clinical implications for the concept of “normal hearing” in the aging population and how older listeners may struggle in different listening environments that are not captured by simple audiometry. Further delineating the important effects of aging will pave the way for earlier clinical diagnosis of hearing disorders and personalized treatments for different forms of age-related hearing loss.

## Introduction

Processing the sounds that occur in everyday life is fundamental to humans’ abilities to interact with each other and with the external world. As we age, our ability to process sounds becomes less robust; older adults with normal audiometric hearing thresholds can understand speech in ideal quiet listening conditions but tend to struggle in more complex listening environments such as those where background noise is present (Gordon-Salant and Fitzgibbons, 1993; Cruickshanks et al., 1998). These complex listening environments degrade the speech signal, making speech understanding more difficult and requiring more effort for correct perception (Gosselin and Gagné, 2011; Pichora-Fuller et al., 2016). This decrease is further exacerbated by age-related temporal-processing deficits, where older adults demonstrated poorer performance on pulse-rate discrimination and gap detection tasks compared to younger adults (Gordon-Salant and Fitzgibbons, 1993; DeVries et al., 2022). It is unclear where the loci of these temporal-processing deficits originate as there are both peripheral and central hypotheses put forth, as well as combined contributions (Willott, 1991; Anderson et al., 2012; Humes et al., 2012).

Aging has long been associated with decreased functions of the auditory system both peripherally and centrally (Roberts and Allen, 2016). One such example is that older adults experience decreased speech comprehension, and this is believed to be associated with neural signal degradation and the subsequent failure to translate these signals into perception (Presacco et al., 2016). Older adults display a decrease in auditory temporal or spectral processing (Harris and Dubno, 2017; Moberly et al., 2018; Xie et al., 2024). More specifically, it has been determined that a deterioration of peripheral auditory structures, which includes peripheral hearing loss in general, as well as the “hidden hearing loss” due to the degeneration of auditory nerve fibers, can augment the auditory processing deficits experienced in aging (Wu et al., 2019; Henry, 2022). Indeed, if spectral- or temporal-processing deficits originate in the periphery, the initial deficits in turn can lead to the altered input of neural signals as information ascends the auditory pathway (Harris et al., 2022).

Paradoxically, auditory stimuli with sound levels above conversational level (e.g., >65 dB SPL) may actually lead to poorer speech perception, especially in the presence of background noise (Molis and Summers, 2003; Summers and Cord, 2007). This decline is known as “rollover.” This phenomenon has been demonstrated in younger and older normal-hearing listeners, as well as older hearing-impaired listeners, and its effects are exacerbated by age and hearing loss (Liu and Kewley-Port, 2007; Jürgensen et al., 2025; Huang et al., 2026). Furthermore, it is likely that temporal features may be linked to poorer speech perception at higher sound levels (Huang et al., 2026). Given that speech itself contains various temporal features, and speech-in-noise stimuli are forms of temporal masking, it is likely that rollover and temporal processing are tightly linked. Given age-related declines in temporal processing, it is expected that even older normal-hearing listeners may be negatively affected by rollover when detecting the important features of speech at high sound levels.

To elucidate the effect of aging on temporal processing in greater detail, we must establish the important relationship between neurophysiology and behavior. Auditory brainstem responses (ABRs) are robust measurements of neural activity in the auditory system spanning from the auditory nerve to the output of the lateral lemniscus and the inferior colliculus (Chiappa and Ropper, 1982; Moller and Jannetta, 1983). The neural activity is represented by key waves within given time windows and each wave represents a different stage of the auditory system. Previous studies have specifically investigated the amplitudes of ABR Wave I (auditory nerve activity), and Wave V (inferior colliculus activity), and found that amplitudes for these waves decreased with age in humans (Jerger and Hall, 1980; Anderson et al., 2021) and animal models (Parthasarathy et al., 2014). Furthermore, aging can also affect the ABR by increasing the latency between waves, possibly due to decreases in conduction velocity (Liberman, 1978; Heil and Irvine, 1997; Bourien et al., 2014). It is thought that deficits of speech intelligibility in noise may be reflected in the characteristics of ABR waveforms in either decreased amplitudes or increased latency for Wave I and V (Burkard and Sims, 2001; Konrad-Martin et al., 2012).

Forward-masking paradigms are often used to measure temporal-processing, where a probe stimulus is masked by a preceding noise stimulus. Thresholds are typically measured by either the minimum possible masker-to-target interval detected or by measuring the minimum sound level in which a probe can be detected at fixed masker-to-target intervals. These studies have been useful in evaluating how temporal-processing deficits arise in the brain by observing changes in the ABR waveform from different forward-masking conditions (Kramer and Teas, 1982; Walton et al., 1999; Hodge et al., 2018). Correlating behavioral thresholds under different forward-masking conditions with changes in the ABR waveform would yield a better understanding of how aging affects the auditory system peripherally and centrally. For example, Mehraei et al. (2017) measured ABR in a forward-masking paradigm and evaluated wave-V latency changes with increasing masker-to-probe intervals in young normal-hearing listeners aged from 20-30 years. In the same listeners, they also measured behavioral forward-masking detection thresholds. The study presented a chirp target stimulus, accounting for the cochlear travelling wave delay (Dau et al., 2000), with and without a preceding masker that was either 35 or 70 dB SPL. The goal of their study was to study these level effects in the context of synaptopathy. They hypothesized that changes in the ABR wave-V latency with forward-masking were related to the loss of low spontaneous-rate (SR), high threshold fibers. This led to larger effects on forward-masking detection thresholds, especially at higher sound levels, and this was supported in their model simulations. They did not, however, apply the modelling results to older listeners experimentally. In summary, while we know that older listeners may potentially be worse at forward-masking tasks and are predicted to have worse temporal-processing abilities at higher sound levels, this remains to be tested explicitly with a chirp target stimulus.

Here we sought to test these questions directly by measuring forward masking behaviorally and electrophysiologically (ABR) to investigate how temporal-processing deficits change across the adult lifespan and with sound level. Using methods similar to Meharei et al. (2017), we investigated the effects of age and level on forward masking using behavioral and electrophysiological approaches. While Mehraei et al. (2017) were interested in hidden hearing loss and thus tested at 35 and 70 dB SPL; the current study was interested in higher intensities where rollover may decrease auditory perception and thus tested at 65 and 80 dB SPL. With increasing age, we hypothesized that there will be increased temporal processing deficits, which will be modulated by the factors of MTI and masker level. As MTI increases, we hypothesized that there would be less forward masking whereas masker level would increase the amount of forward masking because of rollover. We also hypothesized that the ABR data (Wave V) will display similar trends as behavioral detection thresholds, and that the electrophysiology data will be correlated with the behavioral data.

## Materials and Methods

### Listeners

We recruited 33 listeners sampled across decades (20s: N = 8; 30s: N = 5; 40s: N = 4; 50s: N = 4; 60s: N = 5; 70s: N = 7), with clinically normal hearing across the adult lifespan ranging from 20-79 years. Normal hearing was defined as pure-tone thresholds ≤ 25 dB HL (re: ANSI 2018) from 250 to 4000 Hz in the right ear (Fig. 1). Additional criteria included the following: A passing score of ≥ 26 on the Montreal Cognitive Assessment (MoCA; Nasreddine et al. (2005)) and a negative history of neurological disease or middle ear surgery. All procedures were reviewed and approved by the Institutional Review Board (IRB) at the University of Maryland, College Park. Listeners provided informed consent and were compensated for their time.

**Figure 1.**
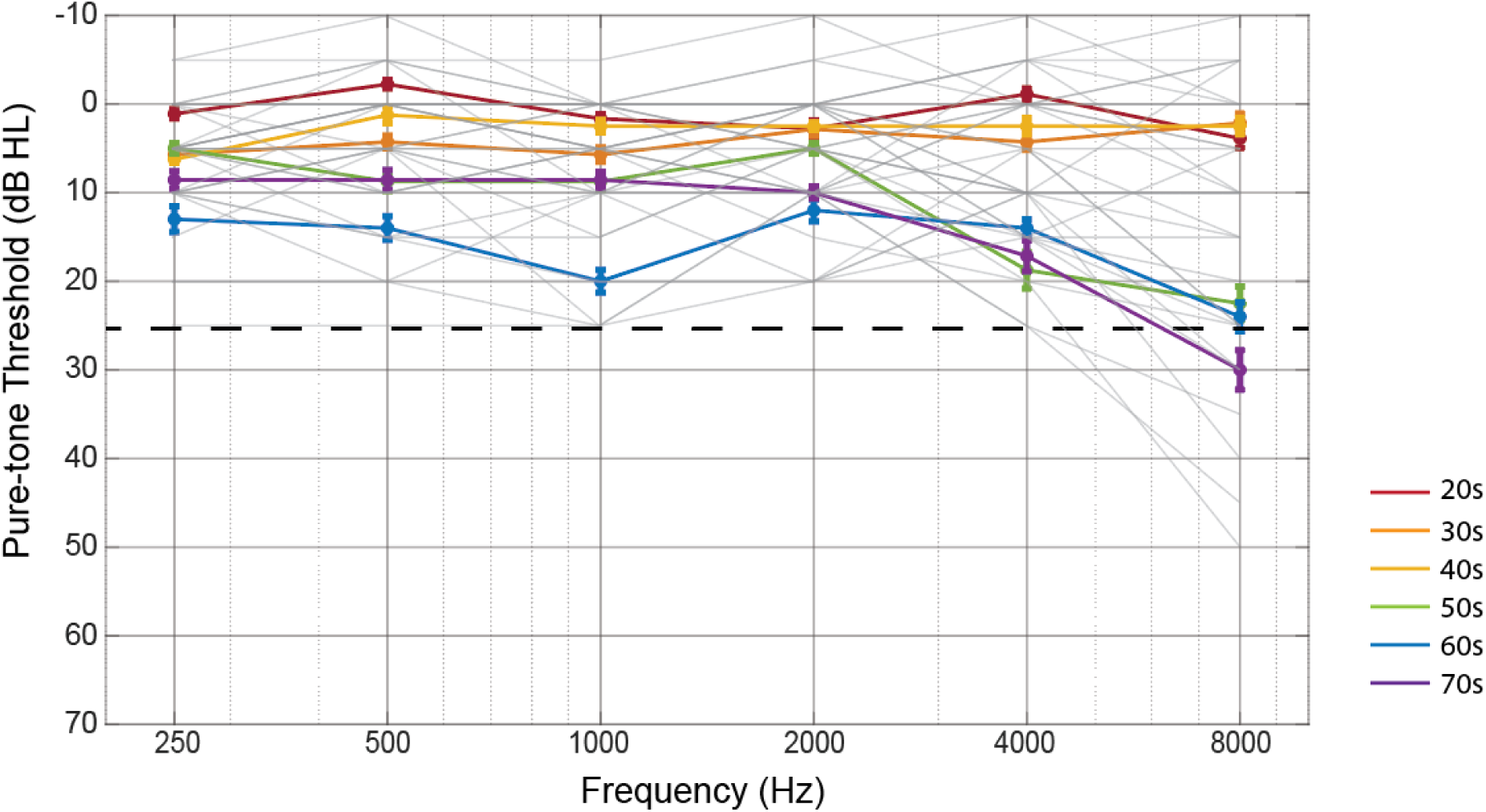
Audiograms for listeners as a function of age. Pure-tone hearing thresholds from 250-8000 Hz for listeners in each decade of life (different colors). Grey lines represent individual hearing thresholds. Dotted line indicates threshold for “normal hearing” at 25 dB HL from 250-4000 Hz. We additionally tested up to 8000 Hz, but this threshold was not considered as criteria for normal hearing.

### Equipment

For the behavioral perception experiment, listeners were seated in a double-walled sound-attenuating booth (IAC Acoustics, North Aurora, IL) in front of a desktop computer where they performed the task. The stimuli were presented monaurally to listeners through a single insert earphone (ER-2, Etymotic, Elk Grove Village, IL). Stimulus presentation was controlled using custom MATLAB scripts (Mathworks, Natick, MA) and amplified to the appropriate sound levels using an amplifier (TDT system 3 HB7, Tucker-Davis Technologies, Alachua, FL). Responses were recorded online to a network drive connected to the same desktop computer with which the listener interacts with the experiment. For the electrophysiology experiment, listeners were seated in a single-walled sound and electrically isolated booth (IAC Acoustics, North Aurora, IL). The stimuli were presented monoaurally to listeners through a single insert earphone (ER-30, Etymotic, Elk Grove Village, IL). Stimulus presentation and event timing were controlled using a custom Presentation script (Neurobehavioral Systems, Berkeley, CA). The BioSemi ActiABR-200 acquisition system (BioSemi B.V., Netherlands) was used to record ABRs. Recordings were obtained with a standard five-electrode vertical montage (Cz active, two forehead offset CMS/DRL electrodes, two earlobe reference electrodes) at a sampling rate of 16,384 Hz.

### Stimuli

The stimuli consisted of two components. First, a 100-ms masking broadband noise was presented at levels of 65 and 80 dB SPL. The noise was temporally shaped with a 20-ms raised-cosine ramp to minimize onset and offset transients. Second, a 15-ms flat-spectrum “synchronized” chirp spanning the frequency range of 40-4000 Hz was used as the target stimulus (Fig. 2A). This chirp is designed to account for the group delay observed in the traveling wave along the cochlea by first presenting low- and then high-frequency components in time (Dau et al., 2000) and thus was intended to maximize the ABR response. The target stimulus followed the masker noise stimulus at MTIs of 5, 50, 100, and 200 ms, which were chosen based on pilot testing.

**Figure 2.**
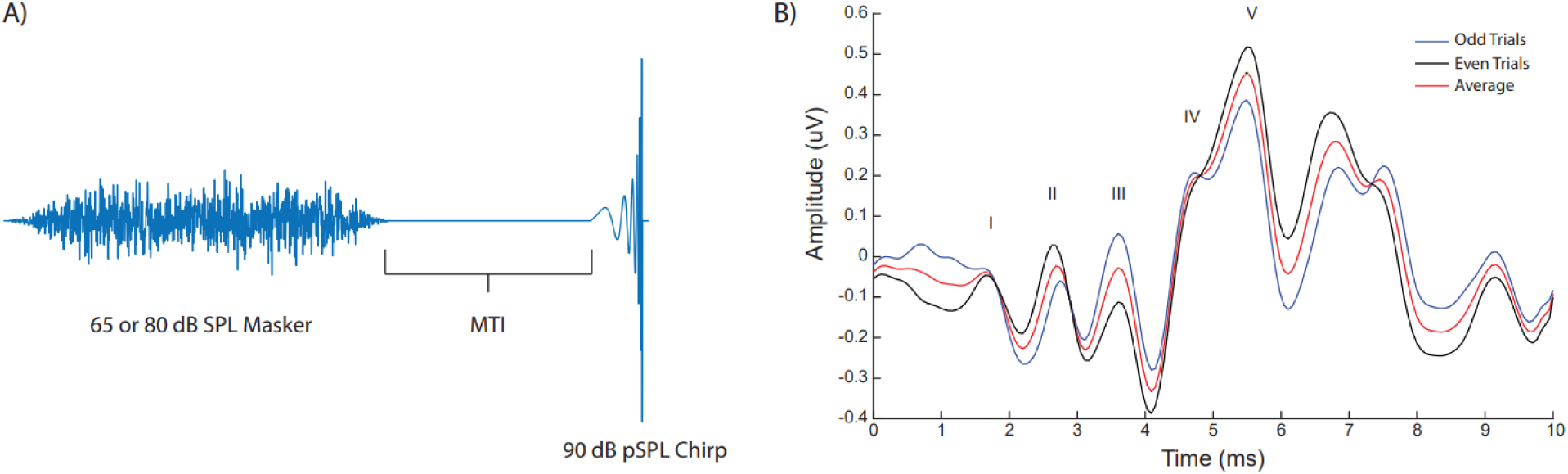
Stimuli and ABR recordings. A) Example of forward masking stimulus, containing a broadband forward masker, followed by one selected MTI, and finally a target chirp stimulus. B) Example ABR demonstrating data collection methodology where averaging occurred across every other trial in “odd” (blue) or “even” (black) trials, as well as the grand average of all trials (red) to ensure replicability in recordings.

### Behavioral forward-masking experiment

On each trial of the behavioral experiment, listeners were presented with a sequence of three sound intervals. The first interval contained only the masker as reference, while the masker followed by the target stimulus was presented randomly in only one of two latter intervals. A silence duration of 300 ms was presented in between each interval to separate them. Thus, this was a 3-interval, 2-alternative forced choice (3I-2AFC) task. Listeners were asked to identify the interval in which the target stimulus was present. Using a 3-down-1-up staircase procedure (Levitt 1971), starting at 65 dB peak dB SPL (pSPL) for the target stimulus, with steps of 5 (reversals 1 and 2), 2 (reversals 3 and 4), and then 1 dB SPL (reversals 5 through 10) to measure the detection threshold for the target stimulus at each MTI combination. The conditions were presented in randomized tracks with 10 reversals allowed and the last 6 reversals were averaged to obtain the detection threshold for the track. Three repetitions of each condition were implemented and were averaged to obtain the final detection threshold for each condition. A total of 8 conditions were tested with combinations of the 4 MTIs x 2 masker Levels.

### Forward-masking ABRs

Forward-masking ABRs were recorded using a Biosemi five-channel EEG. The five-channel configuration included channels CMS and DRL on the forehead, with LA, and RA positioned on the earlobes of the participant with CZ as the reference. ABRs were measured using the same masker and target as in the forward masking behavioral experiment, presented in one rarefaction polarity. However, for the ABRs, the target stimulus level was fixed at 90 dB pSPL to elicit a strong response from the low and high-SR fibers (Dau et al., 2000). We similarly presented the masker and target stimuli at the same four MTIs (5, 50, 100, and 200 ms) with two masker levels (65 and 80 dB SPL), and with the target stimulus alone, yielding a total of nine conditions. A valid trial of the ABR response consisted of a waveform following stimulus presentation which did not exceed a predefined range of ± 50 V for online artifact rejection. A minimum of 2000 valid trials were presented per stimulus condition. The recording session took approximately 2.5 h. The channels were referenced to the average of the two earlobe channels. The recorded data were sampled at 16.384 kHz and imported into MATLAB using custom scripts. The data were then filtered between 0.07 and 2 kHz, to minimize any remaining electrical power line noise at 60 Hz present in the recording. The filtered data were then analyzed at a time epoch from −10 to 10 ms relative to the offset of the chirp in each MTI condition. Replicability of Wave V was ensured by comparing averages of 1000 alternating odd and even trials and Wave V was best identified using the Cz to average earlobe channels; thus, this configuration was used for Wave V amplitude and latency analysis on the combine average of 2000 trials (Fig. 2B). Peak amplitude and latency of Wave-V were identified using visual overlay cursors on a computer screen.

### Forward-masking recovery function

The forward-masking recovery function was modeled following the approach of Jahn et al. (2021). The functions were modeled exponentially using a self-starting asymptotic regression function of the form:

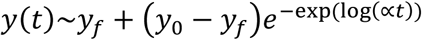

where the measured masker threshold (y) as a function of time starts at y_0_ and decays towards y_f_ at a forward-masking recovery rate α, where larger values indicate faster recovery. The recovery time constant, τ, is derived from 1/α and represents the recovery time constant for forward masking. In other words, τ is the time at which the masker level is reduced to 1/*e ≈* 0.3679 times its initial value (y_0_). We obtained recovery functions for each decade of subjects for analyses.

### Statistical analysis

Mixed-effects logistic regression models were fit to the trial-level data of each stimuli condition, using the *buildmer* package and *lme4* package in *R*.

The model was initially fit for the behavioral data with the dependent variable of the Masked Threshold for the target chirp. This initial model for the behavioral data included fixed factors of MTI and Level, with Age as a continuous variable. Subject was treated as a random effect. A similar approach was taken with the ABR data where the dependent variable was Wave V amplitude or latency. Latency was not further considered due to the fact that in the initial target stimulus condition, the ABR Wave V latency did not significantly correlate with Age, nor were any of the fixed factors of MTI or Level significant. The model was then fit to predict Wave V amplitude for each of the stimuli conditions with fixed factors of MTI, and Level with Age as a continuous variable. Subject was treated as a random effect.

The final mediation model tested whether Wave V amplitude mediated the association of MTI and presentation level with masked threshold, while accounting for age as a continuous covariate. Subject was treated as a random effect.

Model testing was implemented using the *buildmer* function from the *buildmer* package with the binomial family, the default direction for stepwise elimination (the combination of “order” and “backward”), and the default criterion (likelihood-ratio test). Briefly, the *buildmer* function took the initial full model to identify the maximal model that could converge and then performed stepwise elimination to find the best-fitting model for the observed data. The by-participant random intercept was not tested for elimination and was always included in the model.

## Results

### Behavioral forward masking detection thresholds decrease as a function of MTI and increase as a function of age

To obtain a behavioral measure of forward masking, we first tested for thresholds for detection of the chirp stimulus by varying the MTI between the chirp and the preceding masker stimulus (see Methods). We also wanted to confirm previous results that demonstrated that the behavioral detection threshold decreased as MTI increased and investigate how this varies as a function of age. We further tested this with two different masker levels at 65 and 80 dB SPL.

Although here we plotted each decade of age as a separate curve in Fig. 3 to better visualize the data, we used age as a continuous variable in our linear mixed effects model analysis. Overall, we found a main effect of age, indicating that behavioral thresholds in general increase with age at all MTIs for both masker levels. Similar to previous results (Mehraei et al., 2017), we also found a main effect of MTI, indicating that behavioral thresholds decreased as MTI increased. We also found a main effect of Level, where behavioral threshold increased as masker level increased. Finally, there were interaction effects of Age x MTI, and MTI x Level, with steeper decreases at 80 dB than 65 dB SPL (compare Fig 3B with 3A curves). All statistics are described below in Table 1.

**Figure 3.**
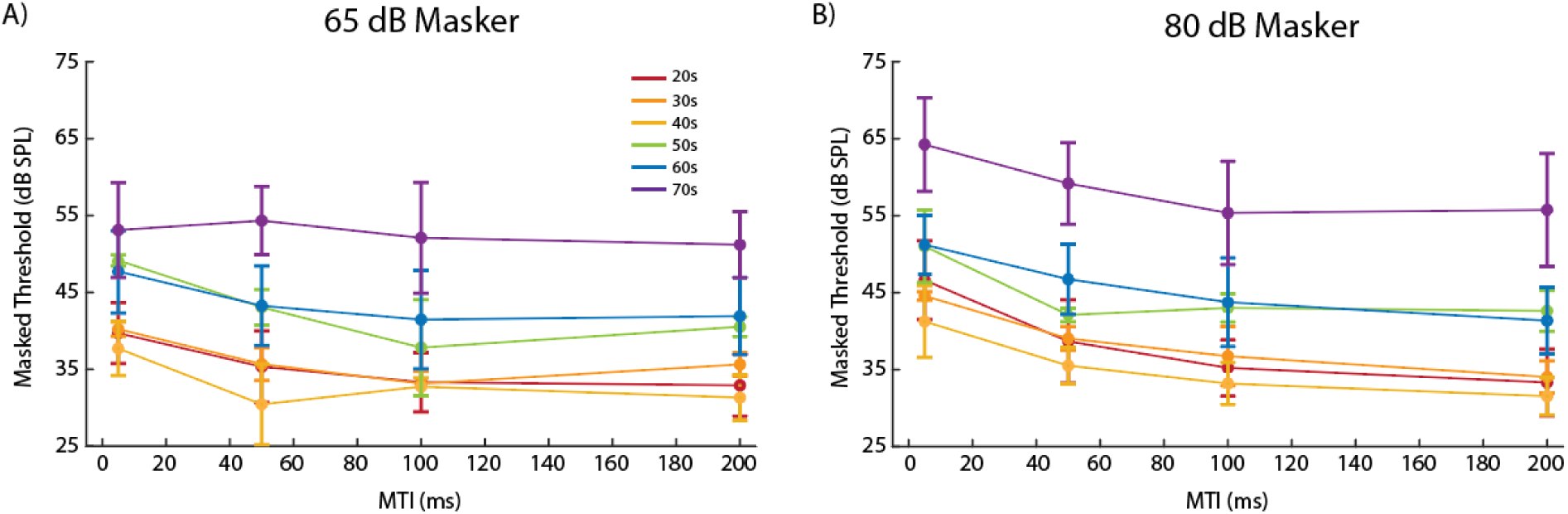
Forward masking thresholds for the behavioral task. A) Masked thresholds as a function of MTI for different decades of age in years with 65 dB SPL masker. B) Same as A but for stimuli with 80 dB SPL masker.

**Table 1.** Results for the best-fitting model for behavior: Masked Threshold = Age + MTI + Level + MTI*Level + Age*MTI + (1 | Subject).

| *Predictors* | *Estimate* | *Std. Error* | *df* | *t value* | *p* |
| --- | --- | --- | --- | --- | --- |
| **(Intercept)** | **42.796** | **1.090** | **31** | **39.278** | **< 0.001** |
| **Age** | **6.112** | **1.092** | **31** | **5.599** | **< 0.001** |
| **MTI** | **-2.639** | **0.202** | **225** | **-13.038** | **< 0.001** |
| **Level** | **1.564** | **0.202** | **225** | **7.728** | **< 0.001** |
| **Age x MTI** | **0.404** | **0.203** | **225** | **1.991** | **0.048** |
| Age x Level | 0.376 | 0.203 | 225 | 1.853 | 0.065 |
| **MTI x Level** | **-0.858** | **0.203** | **225** | **-4.231** | **< 0.001** |
| Age x MTI x Level | 0.139 | 0.203 | 225 | 0.685 | 0.494 |

We used a similar method as Jahn et al., (2021) to fit decaying exponential curves to the behavioral data and obtained τ, which designated the MTI where release from masking would occur. Overall, there were no significant differences between the decades of ages in terms of τ within each masker level, but there was a significant difference between the two levels (t(10) = - 2.845, p = 0.017).

### ABR Wave V amplitudes, but not latency, in response to the target chirp decreases as a function of age

We next wanted to investigate the neurophysiology underlying the main aging effect observed in the behavior data. Using a similar target chirp stimulus as the behavior, but with a fixed pSPL of 90 dB, we presented this stimulus in different masker MTI and Level combinations to the participant in an electrically shielded sound booth while ABRs were collected. We found that presenting the chirp stimulus alone elicited a robust neural response shown in the ABR (Fig. 4A) with a prominent averaged Wave V seen between 5.5 – 6.5 ms, across the decades of ages in our participant pool. Using age as a continuous variable, we found that the Wave V amplitude significantly decreased as age increased (r^2^ = 0.1913, p = 0.0123). However, Wave V latency did not significantly correlate with age (Supplementary Figure 1, r^2^ = 0.047, p = 0.227).

**Figure 4.**
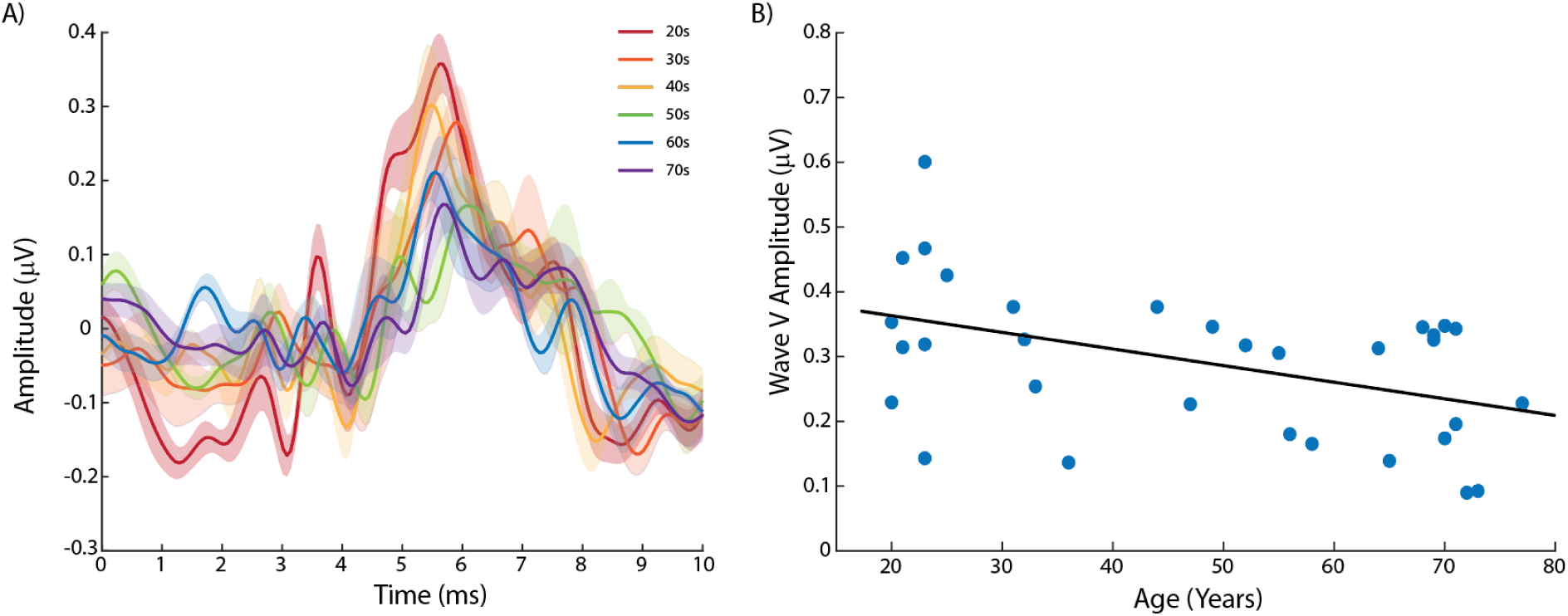
Wave V amplitude in response to target chirp stimulus decreases with age. A) Averaged ABRs elicited by target chirp stimulus alone without preceding masker for each decade of age. Shaded error bars represent ±1 SEM. B) Wave V amplitude as a function of age in years. Solid black line indicates line of best fit.

### ABR Wave V amplitudes, but not latency, are affected by Age, MTI, and Level

Using the same MTI and level conditions as the behavior experiment, we repeated our ABR measurements across the participant pool. For visualization purposes, we plotted the MTI conditions for the two most extreme decades, 20s and 70s, and separated the plots into each masker level (Fig. 5). Similar to our previous data, we found that the overall Wave V amplitude decreased as age increased (compare Fig. 5 A and 5C with 5B and 5D). When analyzing across the participant pool (Table 2), we found that there were significant main effects of Age (β = - 0.053, SE = 0.014, p < 0.001), MTI (β = 0.012, SE = 0.005, p = 0.016), and Level (β = 0.011, SE = 0.005, p = 0.027. There was also a significant interaction of Age x Level (β = -0.01, SE = 0.005, p = 0.036), which suggests that level effects may further drive the differences seen in Wave V amplitude with advancing age.

**Figure 5.**
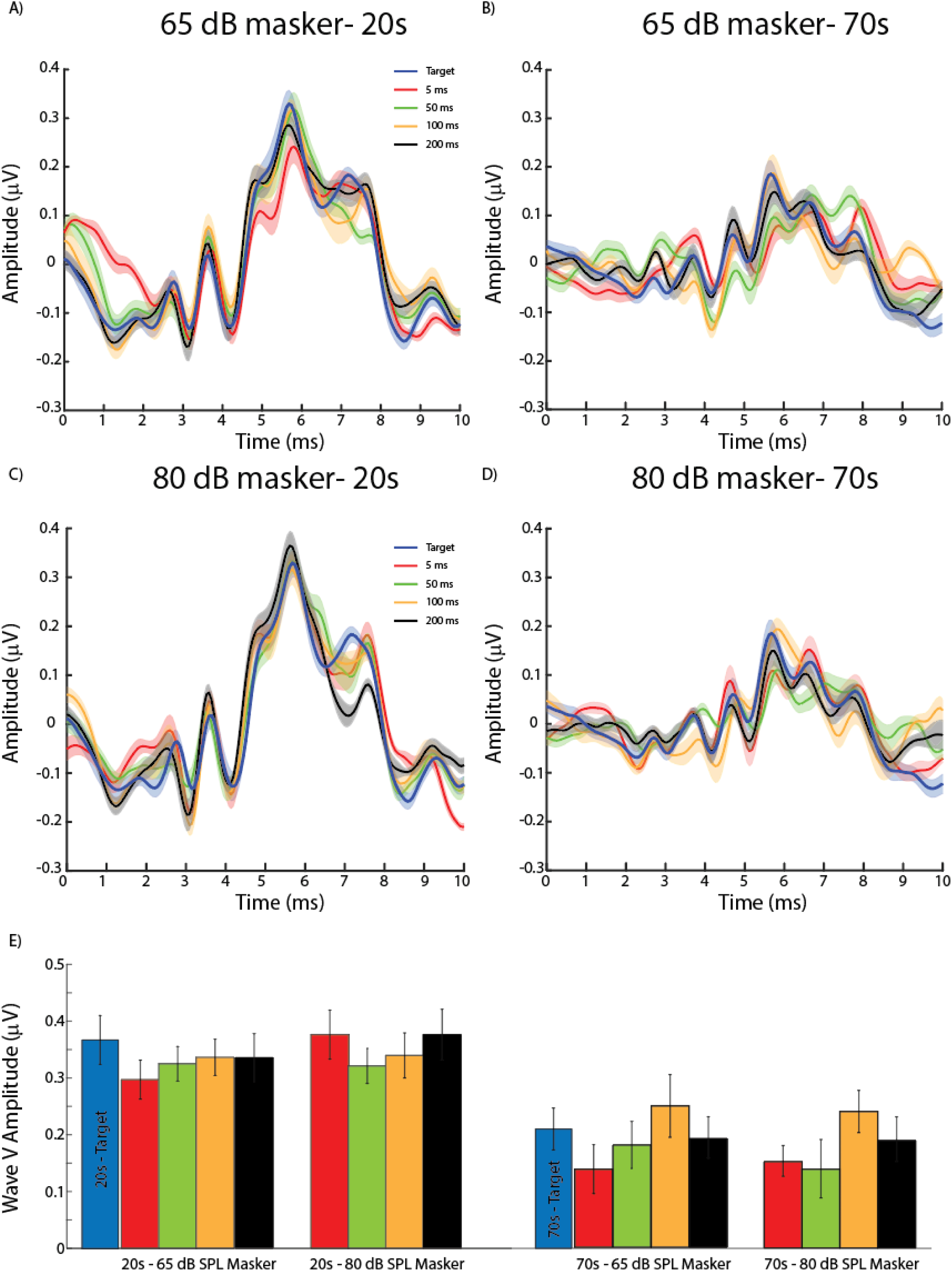
Wave V amplitude differs across age, MTI, and Level. A) Averaged ABR waveform for listeners in 20s decade of age in response to the target chirp stimuli alone (blue) or with a preceding 65 dB SPL masker with MTI of 5 ms (red) 50 ms (green), 100 ms (yellow), and 200 ms (black). B) Same as A for listeners in 70s decade of age. C) Same as A for 80 dB SPL masker. D) Same as B for 80 SPL masker. E) Summary bar graphs comparing conditions plotted in A-D for comparison.

**Table 2.** Results for the best-fitting model for ABR: Wave V Amplitude = Age + MTI + Level + Age*Level + (1 | Subject).

| *Predictors* | *Estimate* | *Std. Error* | *df* | *t value* | *p* |
| --- | --- | --- | --- | --- | --- |
| **(Intercept)** | **0.261** | **0.014** | **31** | **18.635** | **< 0.001** |
| **Age** | **-0.053** | **0.014** | **31** | **-3.768** | **< 0.001** |
| **MTI** | **0.012** | **0.005** | **225** | **2.432** | **0.016** |
| **Level** | **0.011** | **0.005** | **225** | **2.219** | **0.027** |
| Age x MTI | -0.005 | 0.005 | 225 | -0.963 | 0.336 |
| **Age x Level** | **-0.010** | **0.005** | **225** | **-2.108** | **0.036** |
| MTI x Level | -0.006 | 0.005 | 225 | -1.273 | 0.204 |
| Age x MTI x Level | 0.003 | 0.005 | 225 | 0.553 | 0.581 |

Next, we wanted to determine whether these factors had an effect on ABR Wave V latency, as previously observed in Mehraei et al. (2017). When analyzing across the participant pool, we found there were no significant main effects nor significant interactions between Age, Level, and MTI. To ensure that we tested the latency thoroughly, we used both the raw latency value where the peak of Wave V occurred as well as the latency difference between the MTI and the target. Both sets of values could not predicted by Age, Level, or MTI in contrast to Mehraei et al.’s findings. Therefore, for the remainder of the study, we decided to focus on the Wave V amplitude as our primary variable when evaluating forward masking.

### Wave V amplitude modulates forward masking behavior as a function of the interactions between age, MTI and level

Finally, we wanted to determine if we could predict the behavioral results using the neurophysiology from our ABR Wave V data, and asked whether age, MTI, and level all play a role in modulating forward masking. We performed further analysis using a linear mixed effects model to determine if we could predict the behavioral thresholds using our experimental conditions by now including Wave V amplitude in the model (Table 3). The linear mixed effects model revealed main effects of Age, MTI, and Level. There were also significant interactions of Age x MTI, Age x Level, and MTI x Level. Finally, there was a three-way significant interaction of Wave V Amplitude x Age x Level, demonstrating the complex nature of how multiple factors are interacting to drive the behavioral output of forward masking effects.

**Table 3.** Results for the best-fitting model for behavior predicted by neurophysiology data: Masked Threshold = Wave V Amp + Age + MTI + Level + Wave V Amp*Age + Wave V Amp*Level + Age*MTI + Age*Level + MTI*Level + Wave V Amp*Age*Level + (1 | Subject).

| **Predictors** | **Estimate** | **Std. Error** | **df** | **t value** | **p** |
| --- | --- | --- | --- | --- | --- |
| **(Intercept)** | **42.701** | **1.115** | **31.514** | **38.284** | **< 0.001** |
| Wave V Amp | 0.566 | 0.331 | 237.742 | 1.712 | 0.088 |
| **Age** | **6.413** | **1.119** | **31.647** | **5.732** | **< 0.001** |
| **MTI** | **-2.677** | **0.203** | **222.403** | **-13.176** | **< 0.001** |
| **Level** | **1.307** | **0.223** | **222.632** | **5.870** | **< 0.001** |
| Wave V Amp x Age | -0.096 | 0.327 | 237.115 | -0.294 | 0.769 |
| Wave V Amp x MTI | 0.002 | 0.238 | 218.394 | 0.008 | 0.994 |
| **Age x MTI** | **0.415** | **0.203** | **222.405** | **2.044** | **0.042** |
| Wave V Amp x Level | 0.181 | 0.235 | 217.418 | 0.771 | 0.442 |
| **Age x Level** | **0.526** | **0.229** | **222.689** | **2.295** | **0.023** |
| **MTI x Level** | **-0.863** | **0.202** | **221.902** | **-4.268** | **<0.001** |
| Wave V Amp x Age x MTI | -0.003 | 0.206 | 218.038 | -0.014 | 0.988 |
| **Wave V Amp x Age x Level** | **-0.482** | **0.206** | **222.208** | **-2.335** | **0.020** |
| Wave V Amp x MTI x Level | -0.127 | 0.235 | 217.904 | -0.538 | 0.591 |
| Age x MTI x Level | 0.104 | 0.237 | 217.071 | 0.441 | 0.660 |
| Wave V Amp x Age x MTI x Level | 0.300 | 0.204 | 217.710 | 1.473 | 0.142 |

We hypothesized that Wave V amplitude would mediate the effect of the Age × Level interaction on masked thresholds, such that age- and level-related differences in neural processing would contribute to the observed behavioral outcomes. We then broke this Wave V Amp x Age x Level interaction down and assessed the differences while isolating the factors of age and level individually.

Using age as a continuous variable, we first found a significant correlation (r^2^ = 0.151, p < 0.001) between behavioral threshold and ABR Wave V amplitude (Fig. 6A), where Wave V amplitude decreased and masked threshold increased as age increased. This was confirmed by performing a multivariate multiple regression where age was significantly correlated with Wave V Amp (β = 0.809, t(263) = 19.28, p< 0.001) and Masked Threshold (β = 0.004, t(263) = 51.55, p < 0.001). Separating the 65 dB SPL and 80 dB SPL masker level data, we then used a paired-sample Hotelling T^2^ test to determine if there are any differences (Fig. 6B). Using Masked Threshold-Wave V Amplitude as a two-dimensional data point, we found that there was a significant difference between the two masker level populations (T^2^ = 83.39, F(2,130) = 41.38, p < 0.001). Therefore, the combined contributions of Age and Level have an effect on the Wave V amplitude, which then gives rise to the observed behavioral masked thresholds in forward masking.

**Figure 6.**
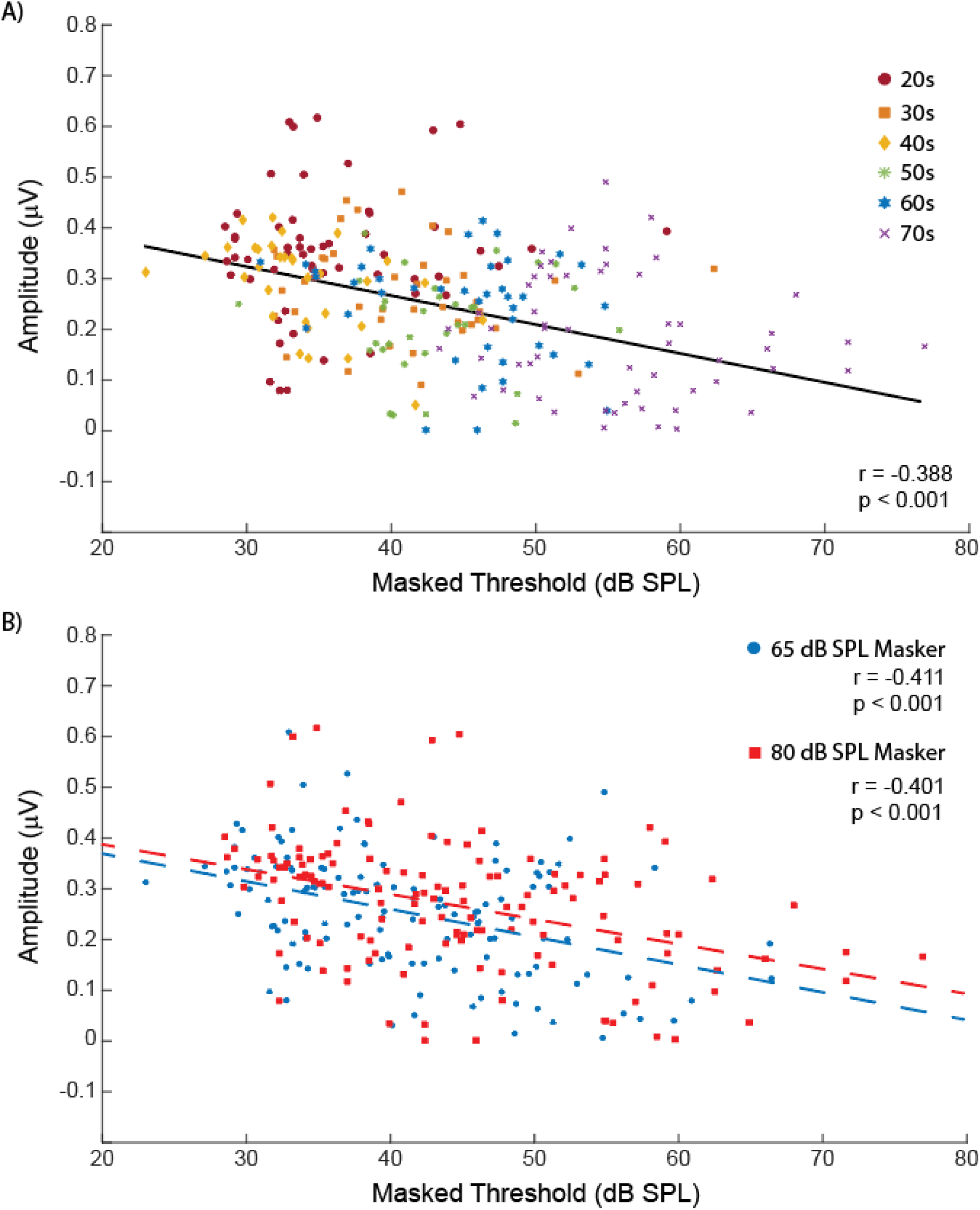
Neural and behavioral response declines as a function of age and is modulated by masker level. A) ABR Wave V amplitude plotted as a function of masked threshold across different age groups. Colors and symbols correspond to each decade of age in years. Solid black line indicates line of best fit. B) Same as A except divided by the two masker levels at 65 dB SPL (blue) and 80 dB SPL (red). Dashed colored lines indicate line of best fit.

## Discussion

### Summary of Results

In this study, we investigated whether the effects of forward masking and level increased as a function of age in audiometrically normal hearing listeners. By testing various MTIs, we were able to determine that release from forward masking in auditory perception occurred at differential values across ages, and the thresholds were overall higher with increased levels. Our electrophysiology results also revealed that ABR Wave V amplitude decreased as a function of age, consistent with behavioral results. Finally, we found a correlation between the electrophysiology and behavioral data, demonstrating that Wave V amplitude and its interactions with age, masker level, and MTI, were predictive of the behavioral thresholds in our forward masking task.

### Forward-masking detection thresholds and aging

Our results demonstrated that forward-masking detection thresholds decreased as MTI increased, but the forward-masking effect appeared to be more prominent with the 80 dB SPL masker (Fig. 3). In general, masked thresholds increased with increasing age for all MTIs and did not converge even at 200 ms MTI across ages (i.e., curves in each decade were parallel). These results combined showed that Age and Level had larger contributions (than MTI) to the observed differences in the masked threshold, and every decade experienced forward masking similarly with one exception. Interestingly, for 65 dB SPL masker, the 70s decade curve (Fig. 3A purple) was relatively flat across MTIs, demonstrating that MTI may not have had any effect on forward masking at that age group. The decade separated data also seemed to suggest that there are three distinct stages of aging as 20-40s decades shared similar thresholds, 50-60s shared similar thresholds, and finally the 70s decade is a group on its own.

Our behavioral findings here were similar to the results found in Mehraei et al. (2017) with their listeners. In that study, they only recruited normal-hearing listeners aged 20-30, which was only limited to one decade of participants in our dataset. The interesting finding here is that our forward masking threshold increased with age (Fig. 3), as we tested across decades up to 79 years of age. Our data in this study demonstrated that temporal masking is greater in older listeners, and they require louder targets to release from the energetic masking from the preceding stimuli. These results are consistent with previous findings in the literature, where older normal-hearing listeners were more greatly affected by forward masking demonstrated by their ABRs (Walton et al., 1999; Heidari et al., 2023). One of the other effects found in our behavioral data here was that detection thresholds increased with increased masker level (compare Fig. 3A and 3B). In contrast to Mehraei et al. (2017), where 35 and 70 dB SPL maskers were used in the stimulus, we instead used a 65 dB SPL masker for a “comfortable” and 80 dB SPL masker for a “loud” level. The levels chosen here in this study were to specifically test the effect of “rollover”, which is the paradoxical decrease of perception as presentation level increases past comfortable hearing levels (Molis and Summers, 2003). It was found previously that increasing the sound past a comfortable conversational level (e.g., 65 dB SPL) to increase audibility does not necessarily yield better speech understanding, and instead higher sound levels can decrease performance, especially in noise (Molis and Summers, 2003). Here, we similarly found that masked thresholds were higher with the 80 dB SPL than for the 65 dB SPL masker, which is also similar to the results found in Mehraei et al. (2017) where the 70 dB SPL masker (which is already past 65 dB SPL), had higher thresholds than at 35 dB SPL. These data further demonstrated that masker level plays a prominent role in forward masking. It is likely that rollover underlies age-related forward masking increases with higher sound levels, emphasizing the role of loudness in temporal processing and perception.

Although we did not directly study the effects of aging on temporal processing of speech here, the forward-masking effects with age found here have implications for poorer speech intelligibility and understanding in older individuals, suggesting that these temporal deficits are directly related to poorer speech perception (Gifford et al., 2007; Tinnemore et al., 2024). It is likely that with rollover occurring at an 80 dB SPL masker, the periphery of the auditory system reached a saturation point faster in older individuals (Liberman and Kujawa, 2017; Parthasarathy and Kujawa, 2018), leading to a greater effect of forward masking. These effects may underlie a form of cochlear synaptopathy that goes undetected in routine audiograms where only pure-tone frequencies in quiet are tested. This cochlear synaptopathy that develops with advanced age may have a direct effect on temporal processing of speech (Carcagno and Plack, 2020, 2022) as well as stimulus interactions with level (Dubno et al., 2012; Carcagno and Plack, 2021). Because synaptopathy, rollover, and age-related temporal processing deficits are all intricately linked, future studies should further measure temporal processing with additional sound levels alongside more ecologically valid stimuli such as speech to validate the relationship between these concepts.

### Relating physiology with behavior

Our electrophysiology results here demonstrated the relationship between physiology and behavior, with Wave V amplitude reliably predicting behavioral detection thresholds (Fig. 6). It was also clear that varying factors, including MTI, masker level, and most importantly age, drive this strong relationship between physiology and behavior. It is likely that age has the most prominent effect on the neural activity, with masker level and MTI stimulus interactions to a lesser extent. The effects of MTI may be too subtle to observe in our electrophysiology data, but are sufficient to explain that forward masking does still occur in our behavioral data where different MTIs resulted in different masked thresholds. Therefore, all three factors (Age, MTI, and Level) contribute and modulate the neural activity measured by Wave V amplitude, which transmits their effects to the observed behavior in the forward masking task.

In contrast to Mehraei et al.’s study, we did not find a significant effect of Wave V latency. This may be due to our sample size within each decade, since Mehraei et al. (2017) only collected ABR data within the decade of 20-30 years of age where latency effects may be the most robust. Each decade of age may be relatively consistent in Wave V latency across subjects, but when evaluating across ages, the increased variability and small sampling of subjects within each decade may explain the differences in our data. Although we did not find an effect of latency, this does not discount the hypothesis that the underlying mechanisms which are responsible for the observed forward masking effects are temporal in nature. For example, the decreased ABR Wave V amplitude with increased age may be explained by the desynchronization of auditory nerve fibers in the periphery, which could be due to cochlear synaptopathy with advancing age (Carcagno and Plack, 2020). The timing and type of ANF firing is critical to the coding fidelity of the auditory system. Desynchronization of these signals is detrimental as temporal information is lost, which can potentially explain the perceptual difficulties experienced by older listeners (Anderson et al., 2012; Furman et al., 2013; Anderson and Karawani, 2020; Arora and Maruthy, 2026).

### Limitations and Future Directions

It is difficult to fully assess the peripheral auditory system without invasive electrophysiology. While we were able to obtain robust ABRs in our younger listeners, the presence of ABR Wave I (similar to Wave V) diminished with increased age. Even with the population average of our decade-by-decade analysis, as ages of our participants approached 60-70s (see Fig. 5), a reliable ABR Wave I could not be obtained and thus the amplitude and latency were not further analyzed in this study.

Furthermore, there was not a significant effect of latency in our ABR data as mentioned previously, and this could be explained by the increased variability observed in our participant pool within each decade and across ages. While we chose to use a chirp as the target stimulus to maximize responses, the variability in the latency data may be a result of additional unknown cochlear mechanics in response to the chirp. We observed this when we piloted the study by presenting the same chirp in opposite polarities, which resulted in ABRs that were phase shifted, but only in a subset of our participants. This finding contrasted with previous studies, in which a click target stimulus, either alone or with a masker was used rather than a chirp, yielded more consistent effects on latency (Burkard and Sims, 2001; Anderson et al., 2021; Poe et al., 2024). Future studies may revisit the stimuli parameters used to produce more robust latency effects with alternative maskers or presentation levels of the target stimulus. A systematic investigation of stimulus parameters may help us understand how to completely maximize neural and behavioral responses for forward masking studies in the future.

While we did not explicitly test speech stimuli here, our results have many implications for temporal processing of speech as mentioned above. Future studies should focus on ecologically valid maskers and target stimuli such as presenting multi-talker babble preceding or following a target syllable or word. Finally, our results here demonstrate that clinical audiograms may not necessarily capture the full extent of age-related hearing loss and the “hidden hearing loss” experienced by older adults may require refinement of future diagnosis and treatment in the clinic.

## Ethical approval and informed consent statements

All procedures were reviewed and approved by the Institutional Review Board at the University of Maryland, College Park. All listeners provided informed consent in writing and were compensated for their time.

## Data availability statement

All data for the manuscript are available on Figshare. The link for the data is available from the corresponding author upon request.

---

**Supplementary Figure 1.**
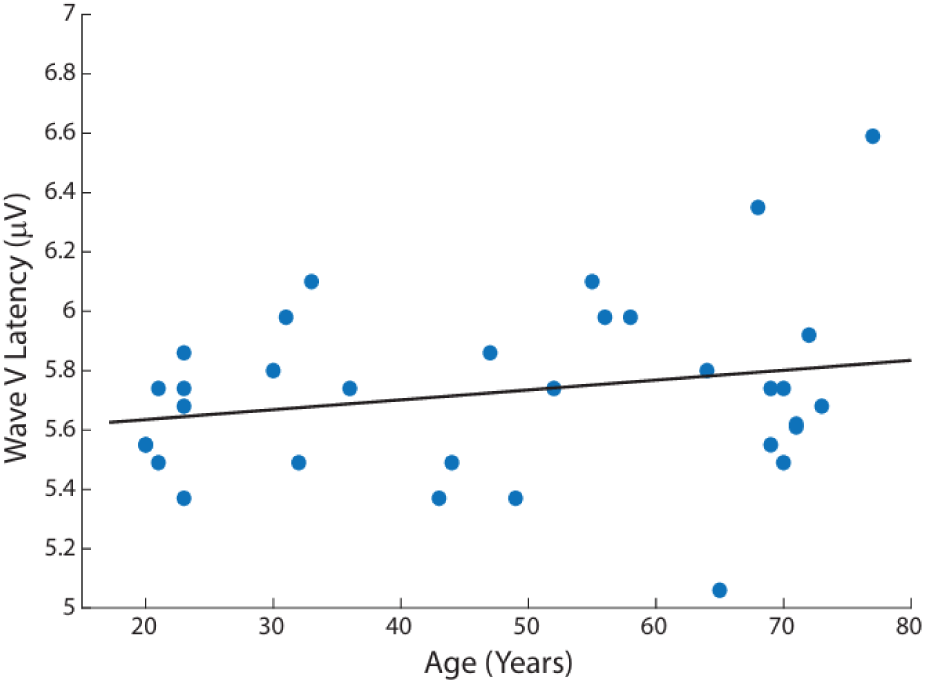
Wave V latency in response to target chirp stimulus does not correlate with age. Wave V latency as a function of age in years. Solid black line indicates line of best fit.

