## Supplementary figures and images for "Auditory brainstem response amplitude predicts age-related forward masking temporal-processing deficits in normal-hearing listeners"

### Supplemental Figure 1

#### Supplementary Figure 1

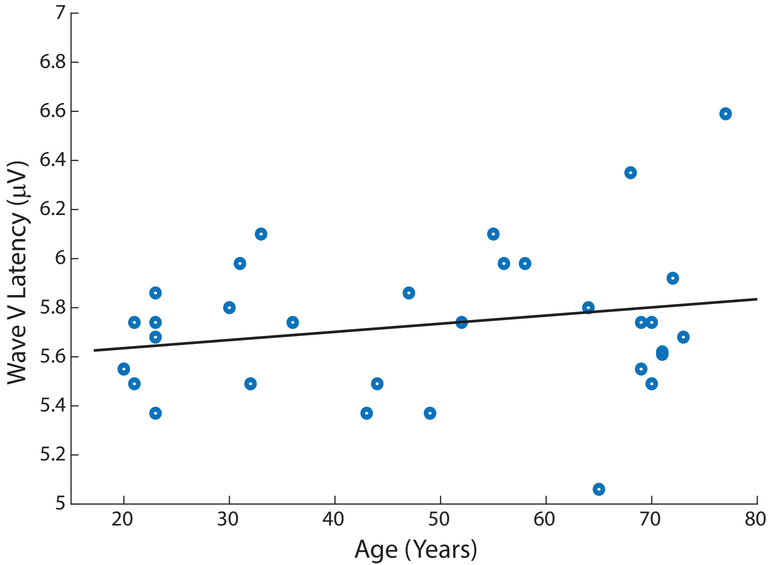
